# K-means Based Unsupervised Feature Selection to Prioritize Biomarkers of Different Disease Clinical Phases

**DOI:** 10.1101/2020.04.21.052704

**Authors:** Xue Jiang, Weidi Wang, Jing Xu, Zhen Wang, Guan Ning Lin

## Abstract

Huntington’s disease is caused by a single gene mutation, which is potentially a good model for development of biomarkers corresponding to different disease phase and clinical phenotypes. Hypothesis-driven and omics discovery approaches have not yet identified effective candidate biomarkers in HD. So, it is urgent to develop engagement and disease-phase specific biomarkers. The advanced sequencing technology makes it possible to develop data-driven methods for biomarkers discovery. Therefore, in this study, we designed k-means based unsupervised feature selection (KFS) method to prioritize biomarkers of different disease clinical phases. KFS first conducts k-means clustering on the samples with gene expression data, then it conducts feature selection based on the feature selection matrix to prioritize biomarkers of different samples. By conducting alternative iteration of clustering and feature selection to screen key genes which corresponding to the complex clinical phenotypes of different disease phases. Further gene ontology and enrichment analysis highlight potential molecular mechanisms of HD. Our experimental analyses have uncovered new disease-related genes and disease-associated pathways, which in turn have provided insight into the molecular mechanisms during the disease progression.

## I. Introduction

Huntington’s disease (HD) is caused by a single genetic insert structure mutation (CAG repeat expansion) in the huntingtin (HTT) gene on chromosome 4 that codes for polyglutamine in the huntingtin protein. Mutant huntingtin can affect cellular metabolism, mitochondrial function, which lead to the production of abnormal metabolites and markers of oxidative stress [1]. Besides, the mutant huntingtin protein can enter the nucleus and alter gene transcription [2], and produce many abnormal effects in cells, such as abnormalities in cellular proteostasis mechanisms [3]. Earliest steps in the pathogenic cascade of HD include misfolding of huntingtin to a beta-sheet structure, and post-translational alterations, such as cleavage or altered phosphorylation.

Age of onset and rate of progression of HD are both likely to be influenced by environmental and genetic modifies [4], [5]. Clinically, motor, cognitive, and emotional function are severely affected during the disease progression. The severity of symptoms are positive correlated with age and CAG repeat length. Now days, Huntington’s disease can be predicted by genetic testing. However, the molecular mechanisms under complex clinical phenotypes still have many unknown. Many pathogenic mechanisms have been hypothesized for HD, promoting the biomarker development. Hypothesis-driven studies mainly focus on functional correlates and neurobiological underpinnings of detectable changes that already have been reported, such as immune activation [6]–[8], transcriptional dysregulation [9] and cholesterol biosynthesis. Omics discovery approaches have not yet identified good candidate biomarkers in HD [10]. So, there is a great need exists for target engagement and treatment-specific biomarkers. However, at present, there is no effective technology to treat for the disease. The continued development of functional, neurochemical and other biomarkers raises hopes that these biomarkers might be useful for future trials of disease-modifying therapeutics to delay the onset and slow the progression of HD. The advances in biomarker discovery also could herald new era of personalized preventive therapy.

The development of high-throughput sequencing technology enables researchers to prioritize disease-association biomarkers with data-driven approaches, such as machine learning methods and deep learning methods [11]–[16]. Correlating those biomarkers to clinical trials can promoting the understanding of physiopathologic mechanisms under the abnormal behavior. It has been reported that cognitive, motor, and emotional functions have been gradually affected by the Huntington’s disease. Cognitive impairments usually emerge years before diagnosis of HD. Motor features are most prominent with adult-onset or late-onset HD, begin early in the course of the disease, and give HD its characteristic clinical appearance. In addition, the emotional features are more variable than are the motor or cognitive features. To decode the molecular mechanisms under those complicate phenotypes, it is better to develop new methods and concepts to prioritize biomarkers of different disease phases and clinical phenotypes with big omics data.

The advanced high-throughput sequencing technology enables researchers to prioritize disease biomarkers in genomewide level, including transcription factor binding site, hypersensitive sites, expression trait locus, and various genetic mutations. These candidate biomarkers can be further intergrated into a molecular interaction network to form a system understanding of the disease development [17], [18]. To select biomarkers of different disease phases and clinical phenotypes, we designed k-means based unsupervised feature selection method (KFS). KFS first conducts k-means clustering on the samples with gene expression data, then it conducts feature selection based on the feature selection matrix to prioritize biomarkers of different samples. By conducting alternative iteration of clustering and feature selection to screen key genes which corresponding to the complex clinical phenotypes of different disease phases. Biomarkers for different severity of disease symptoms are selected out. We also integrated top ranking genes to get a robust disease-related biomarkers. Finally, by performing gene ontology annotation, protein-protein interaction analysis, and functional enrichment analysis, those marker could provided insight into the molecular mechanisms during the disease progression.

The rest of this paper is organized as follows: In Section 2, we present the gene expression data used in this study, and present the proposed KFS model in detail. In Section 3, we present the experimental results of differentially expressed genes obtained by KFS. The single cell and tissue-specific gene expression are analyzed. The function annotation and pathways involved in disease development are further analyzed and reported. In Section 4, conclusions are presented.

## II. Materials and Methods

### A. Gene Expression Data

We downloaded gene expression data from http://www.hdinhd.org. The RNA-seq expression data were obtained from the striatum tissue of Huntington’s disease mice model. The age of experimental mice is 6-month-old. There are 6 genotypes for the gene expression data, including poly Q20, poly Q80, poly Q92, poly Q110, poly Q140, and poly Q175. There are 8 samples for each genotype. Totally, there are 48 samples. It should be noted that poly Q20 is normal mice, and mice with other genotypes are disease ones. The severity of disease symptoms is increasing with the repeat number of CAG [20], [21]. The detailed information of the dataset are illustrated in Table I.

**TABLE I.** Experimental data description.

|  |  |  |  |  |  |  |
| --- | --- | --- | --- | --- | --- | --- |
| Species | mouse |  |  |  |  |  |
| Tissue | striatum |  |  |  |  |  |
| Age | 6-month-old |  |  |  |  |  |
| Genotype | poly Q20 | poly Q80 | poly Q92 | poly Q111 | poly Q140 | poly Q175 |

**TABLE II.**
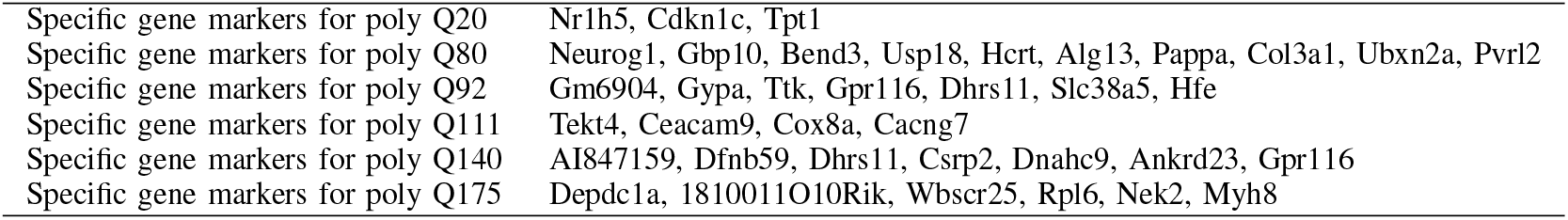
SPECIFIC GENE MARKER FOR DIFFERENT GENOTYPES.

### B. K-means Based Unsupervised Feature Selection

The gene expression data is denoted as *X* = [*x*_*ij*_]_*n*×*m*_, where *x*_*ij*_ represents the expression level of gene *j* in sample *i*. *H* denotes a vector function that can be represent by *H* : *X* → *R*^*c*^. Each sample *x*_*i*_ corresponding to a *H*_*i*_. Let 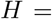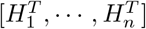 denotes a *n* × *c* cluster indicator matrix, while *c* is the cluster number. *C* = [*c*_*ij*_]_*c*×*m*_ is the cluster center matrix. *F* = [*f*_*ij*_]_*m*×*c*_ is feature selection matrix. Let *A* = [*a*_*ij*_]. We define 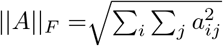, and the *l*_2,1_ of matrix *A* is 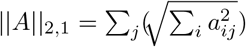.

We hope to conduct clustering of different genotypes, prioritizing key genes that can distinguish different genotypes, thus to understand the molecular mechanisms behind in different onset age caused by the CAG repeat length. In this study, we designed k-means based unsupervised feature selection to conduct clustering and feature selection through an alternating iterations. Since the simple and effective of k-means clustering method, we designed the unsupervised feature selection method to screen key genes that distinguish different samples. K-means is used to capture the inner structure of the data during the clustering process. As not all the features contribute to the sample clustering, we conduct *l*_2,1_ constraint to the feature selection learning process, making the columns sparse. KFS can be modeled as the following function

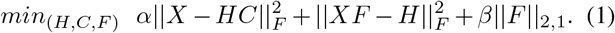

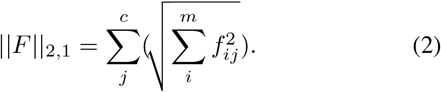

Here, *α* and *β* are hyper-parameters, which balance the importance of the clustering results and sparse feature selection results. For the Eq. 1, we designed an iterative updating algorithm that alternatively updates *F* and *H*. The detailed solving processes are shown below:

Step 1. Initialize the feature selection matrix *F* with random numbers in [0, 1], and select *c* samples in *X* to construct cluster matrix *C*.

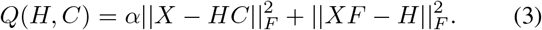

The elements in cluster indicator matrix must be larger than 0. Let *z*_*ij*_ be the Lagrange multiplier that constraint *h*_*ij*_ ≥ 0, the Lagrange function can be written as

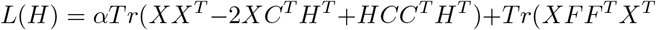

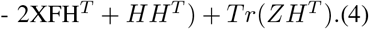

Meanwhile *Z* = [*z*_*ij*_]. The derivation of *H* is

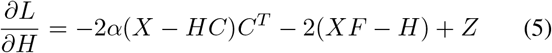

Based on the KKT condition *z*_*ij*_*h*_*ij*_ = 0, we can get

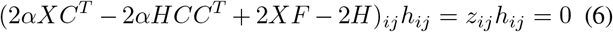

Eq. (6) can be written as

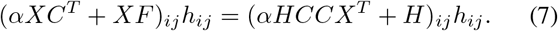

Then, we can get the update role for *H*

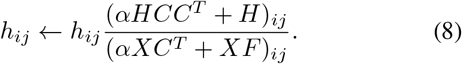

The final label of sample *x*_*i*_ is

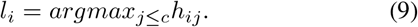

Then we can update the cluster indicator matrix 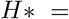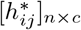 according to Eq. 10.

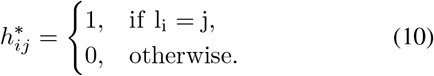

From 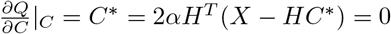, we can get

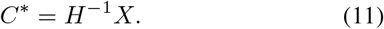

Step 3. According to the last two terms in Eq. 1, we can get the solution for feature selection matrix *F* * by solving the Eq. 12.

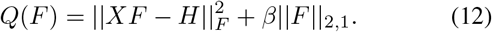

Let *ψ*_*ij*_ be the Lagrange multiplier that constraint *f*_*ij*_ ≥ 0. The Lagrange function can be written as

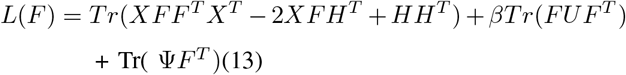

Here, 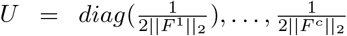 is an Auxiliary matrix, and *F*^*i*^ is denoted as the *i*_*t*_*h* column of matrix *F*, Ψ = [*ψ*_*ij*_].

The derivation of *F* is

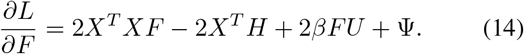

According to the KKT condition *ψ*_*ij*_*f*_*ij*_ = 0, we can get

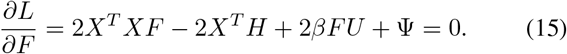

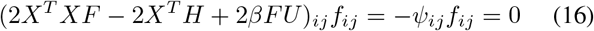

Equation (16) can be written as

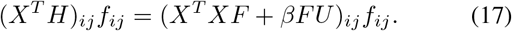

Then, we can get the update role for *F*

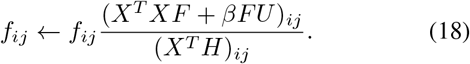

Step 4. Conduct loop iteration of Step 2 until the objective function of Eq. 3 converges. At this point we can get cluster indicator matrix *H*\* and cluster center matrix *C*\*. We set 100 times iteration.

Step 5. Conduct loop iteration of Step 3 until the objective function of Eq. 12 converges. Then we get feature selection matrix *F* *.

Step 6. Conduct loop iteration of Step 4 and Step 5 until the objective function of Eq. 1 converges. We set 10 times iteration for the experiments in this study. In each iteration, the low ranking 500 columns were deleted and the gene expression matrix were modified correspondingly.

Step 7. Based on rank-product method [19], we calculates the element fluctuation of each row in the feature selection matrix. The elements of *i* – *th* row fluctuate significantly, indicating that the corresponding feature *i* (i.e. gene *i*) has a strong ability to distinguish samples of different categories. Sorting the rank-product value of each row in the feature selection matrix in descending order, we select the features that identical to the high ranking row numbers, and remove the features that identical to the row numbers at the bottom of ranking list. Update gene expression matrix *X*.

Step 8. Repeat aforementioned Step 1 to Step 7, we get the final cluster indicator matrix *H*\*, cluster center matrix *C*\*, and the feature selection matrix *F* *.

Let function *top*(*v*_*s*_) represents the *s* greater elements of vector *v*.

Since greater elements in *F*^*j*^ contribute more to specific identification of category *j*, the genes, whose column number in the gene expression matrix is equal to the row number of the greater element in the feature selection matrix, are seen as key features of category *j*, i.e. the genes that are severely affected under this condition. In this study, we use *key*_*j*_ to denote the key gene set of category *j*.

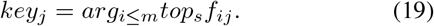

The detailed process of KFS is shown below.

To ensure the convergence of Equation (1), we first solve cluster indicator matrix and cluster center matrix, then we solve feature selection matrix. In this study, we set *α* = 0.8, *β* = 0.2. In each interaction, we remove low ranking 500 genes to modify the gene expression data for next iteration. In this study, we conduct 10 iterations to end the process.

## III. Results

We used RNA-seq expression of Huntington’s disease mice model under 6-month-old with genotypes of poly Q20, poly Q80, poly Q92, poly Q111, poly Q140, and poly Q175 to conduct experiments. Totally, there are 23351 genes. Firstly, we filtered out genes that are not express in any samples. After the filtering, there are 20772 genes left. Then, we scaled the gene expression with minmax scalar. we computed the means and variances of all genes in the samples, and ranked the genes in descending order. The means and variances of gene expression in all samples are shown in Fig. 2 and Fig. 3. To improve the credibility of biomarkers, top 5000 genes were selected for next analysis.

**Fig. 1.**
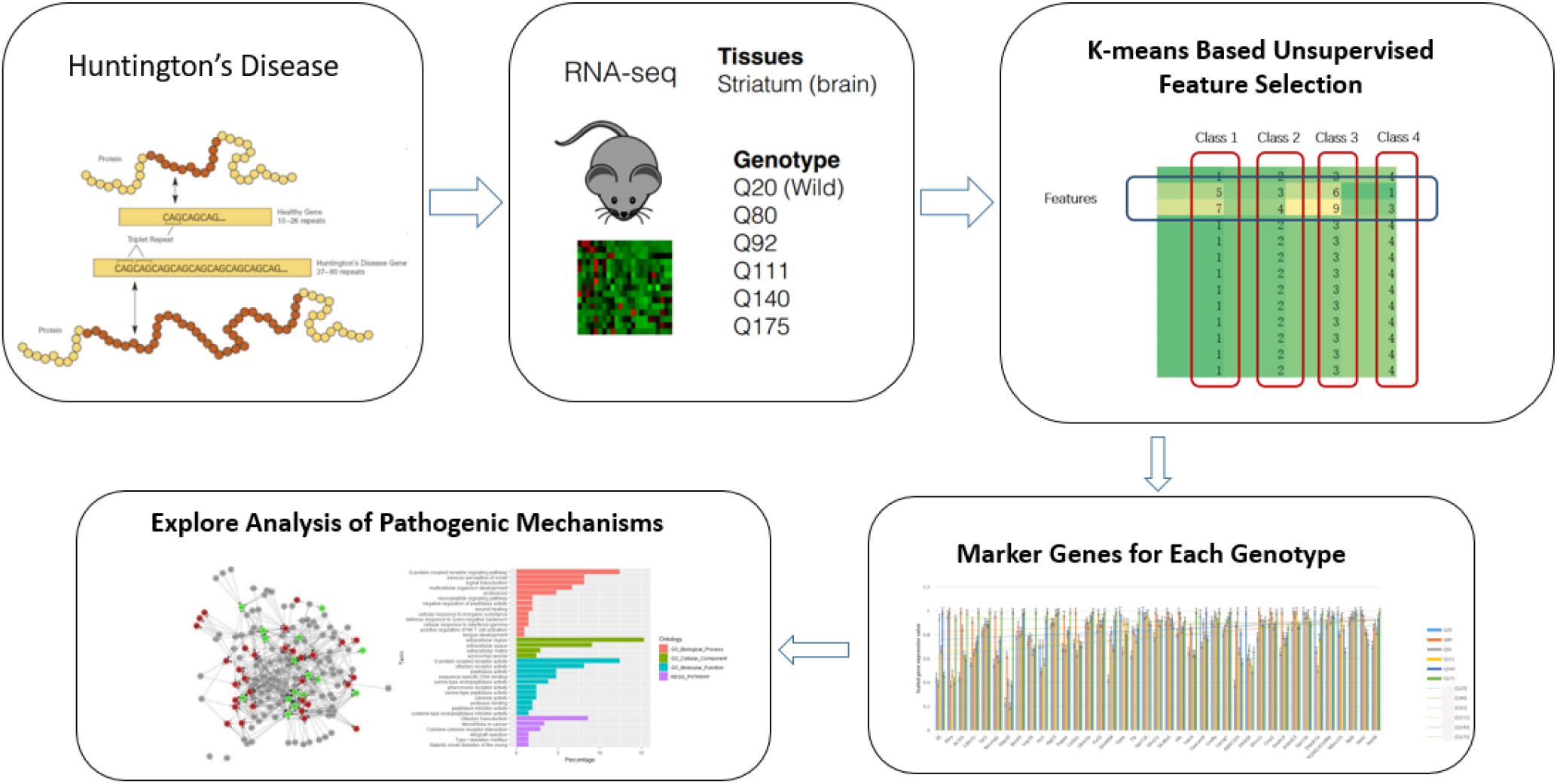
Flowchart of k-means based unsupervised feature selection to prioritize biomarkers of different Huntington’s disease Clinical Phases.

**Fig. 2.**
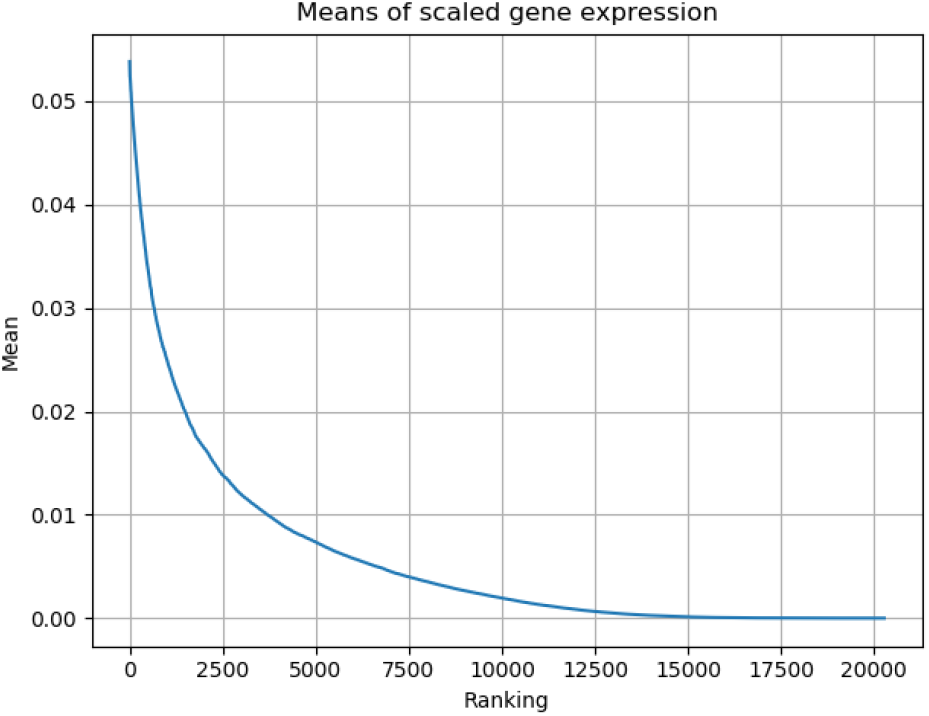
Ranking of the means of geneexpression values in all samples.

**Fig. 3.**
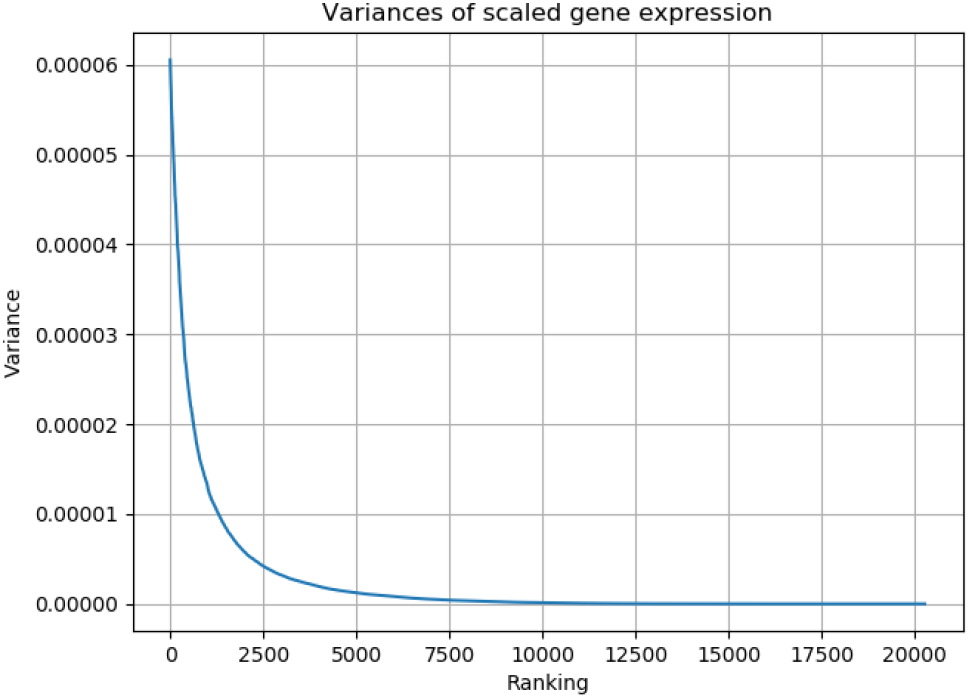
Ranking of the variances of geneexpression values in all samples.

**Fig. 4.**
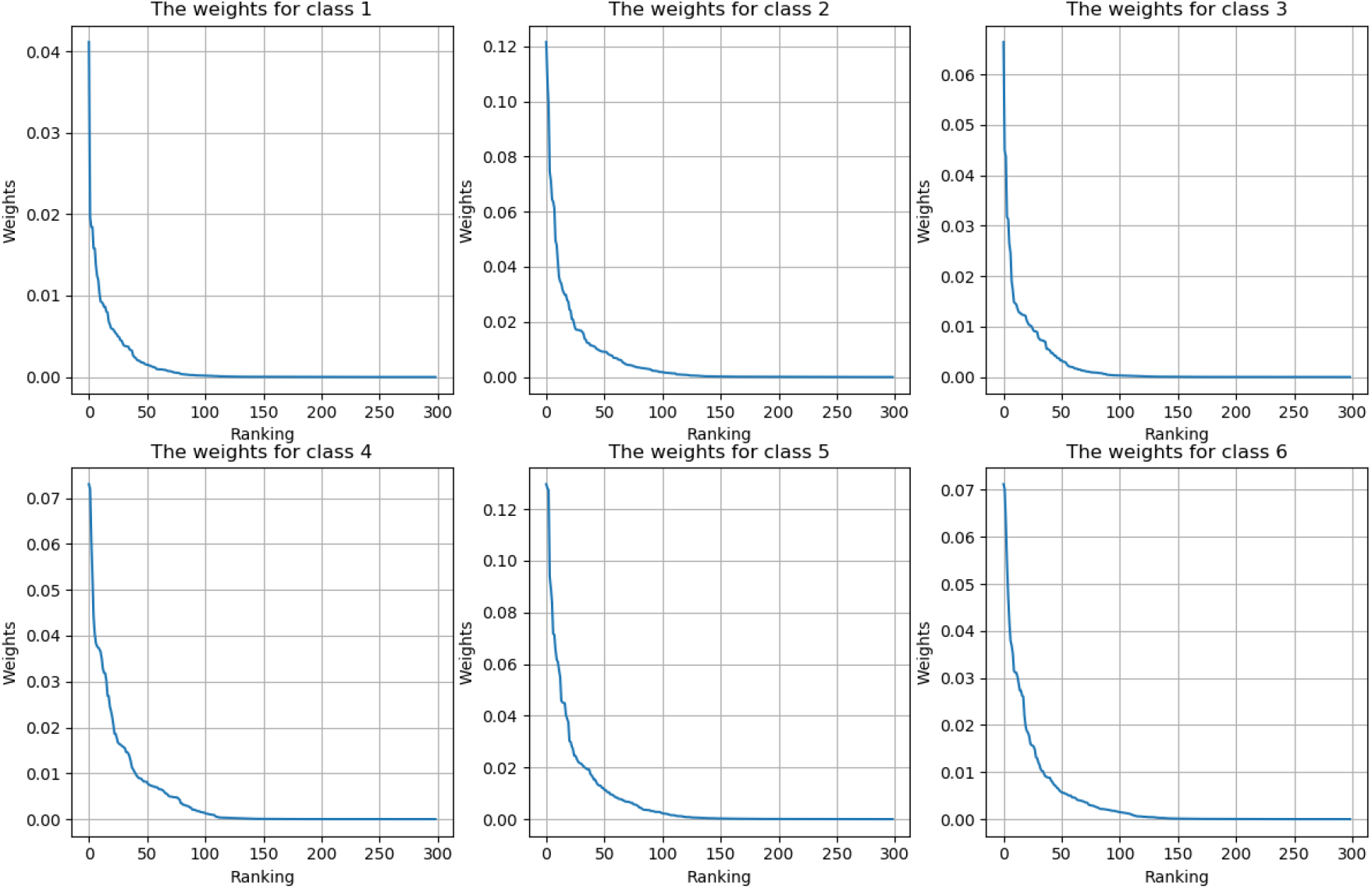
Gene rankings according to the elements in feature selection matrix.

### A. Performance of KFS

We used samples with genotypes of poly Q20, poly Q80, poly Q92, poly Q111, poly Q140, poly Q175 to conduct KFS. Each genotype is seen to be a category. In the 10 iterations of KFS, we got 10 feature selection matrixs. We ranked genes in descending order according to the elements in each column of the feature selection matrix. The rankings of elements in feature selection matrix for the six classes are shown in Fig. 5.

**Fig. 5.**
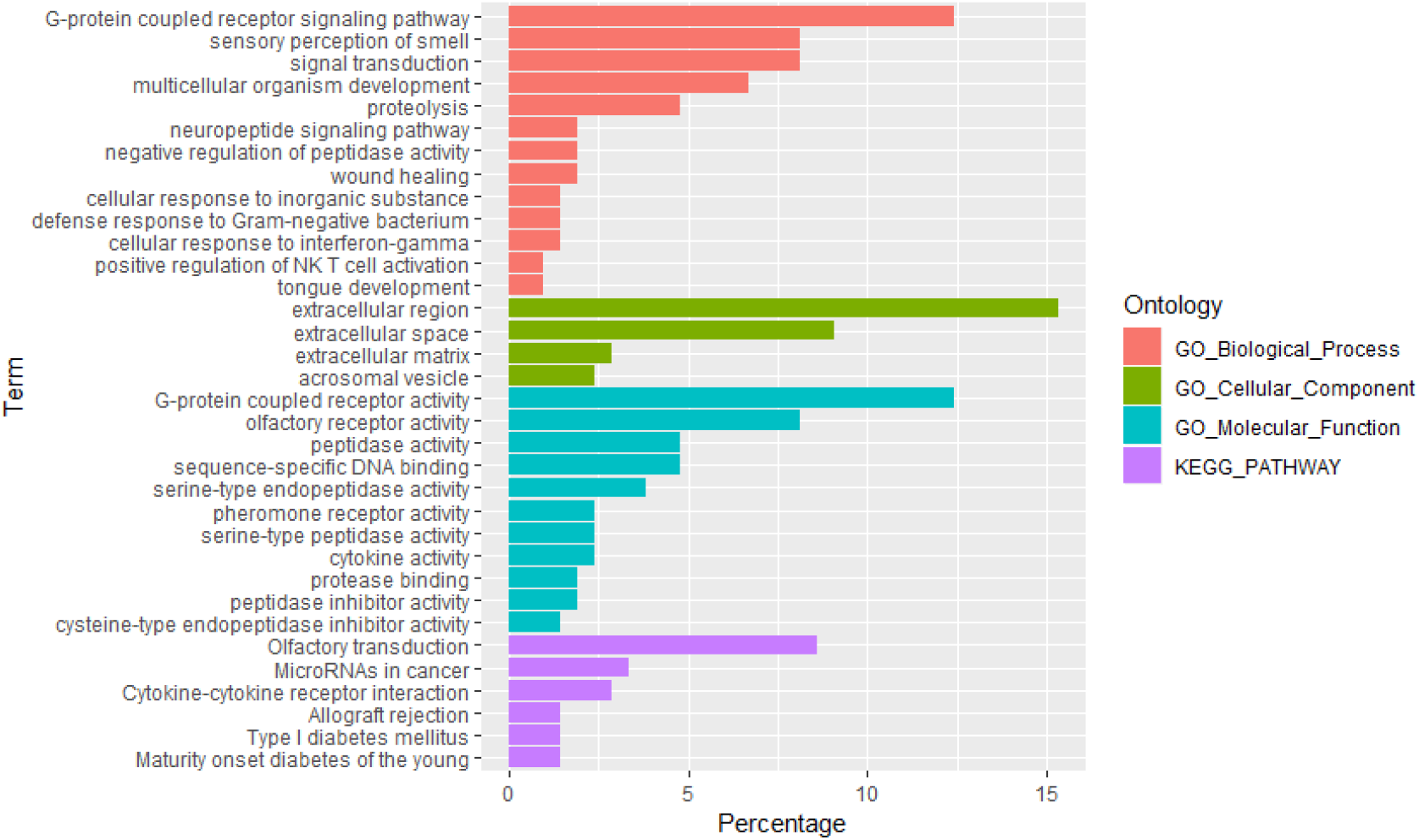
The enrichment terms of 227 disease candidate genes prioritized by KFS.

Through the intersection of top 1000 genes of each iteration. We got final key gene set for each category. KFS selected 3 specific gene markers for poly Q20, 10 specific gene markers for poly Q80, 7 specific gene markers for poly Q92, 4 specific gene markers for poly Q111, 7 specific gene markers for poly Q140, and 6 specific gene markers for poly Q175. Those gene markers are shown in

**Algorithm 1:**
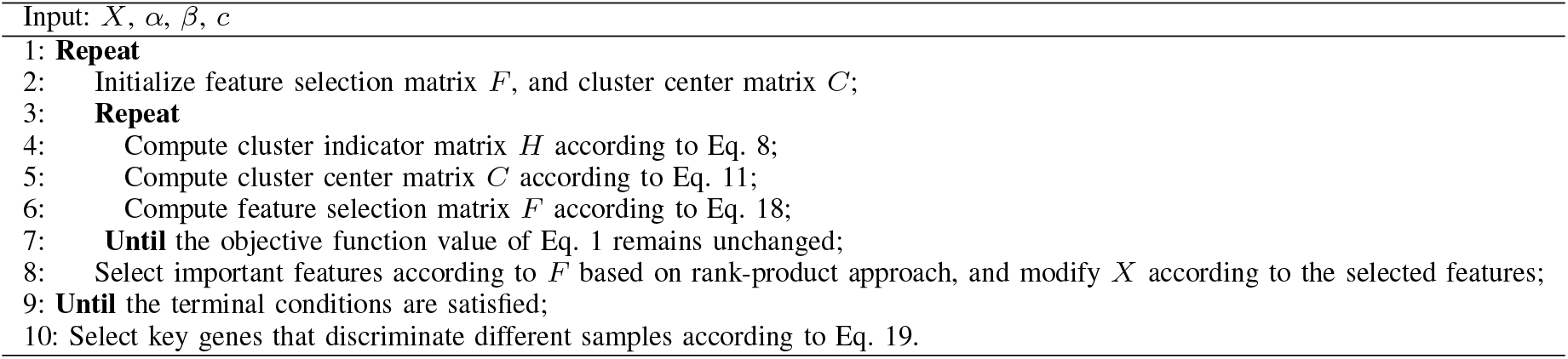
KFS

We also combined top 1000 genes of each iteration to get a robust gene set. KFS prioritized 227 disease candidate genes totally. We used DAVID [22] to conduct enrichment analysis of the 227 disease candidate genes. The enrichment terms of the 227 genes are shown in Fig. 5. From the enrichment analysis, we can know that G-protein coupled receptor signaling pathway and neuropeptide signaling pathway are seriously affected by the excessive CAG duplication in HTT gene.

We used the STRING (https://string-db.org) to examine protein protein interactions among the 227 genes. Two hub genes, IL5 and IL5ra were selected. IL5 and IL5ra are all related with extracellular space, and mainly correlated to biological processes of immune response and inflammatory response.

We further demonstrated the scaled expression values of those 39 marker genes under different genotypes, see Fig. 6. From Fig. 6, we can clearly know that the expression values of those genes were up-regulated in disease mice compared with that in normal mice. The trend lines of gene expression value of different phenotypes indicate that the gene expression levels of those marker genes are more stable in disease mice compared with that in mice of genotype poly Q20. We further used the PsyMuKB (http://www.psymukb.net/) to annotated the 39 marker genes, meanwhile Tpt1, Neurog1, Alg13, Col3a1 and Ttk are human essential genes.

**Fig. 6.**
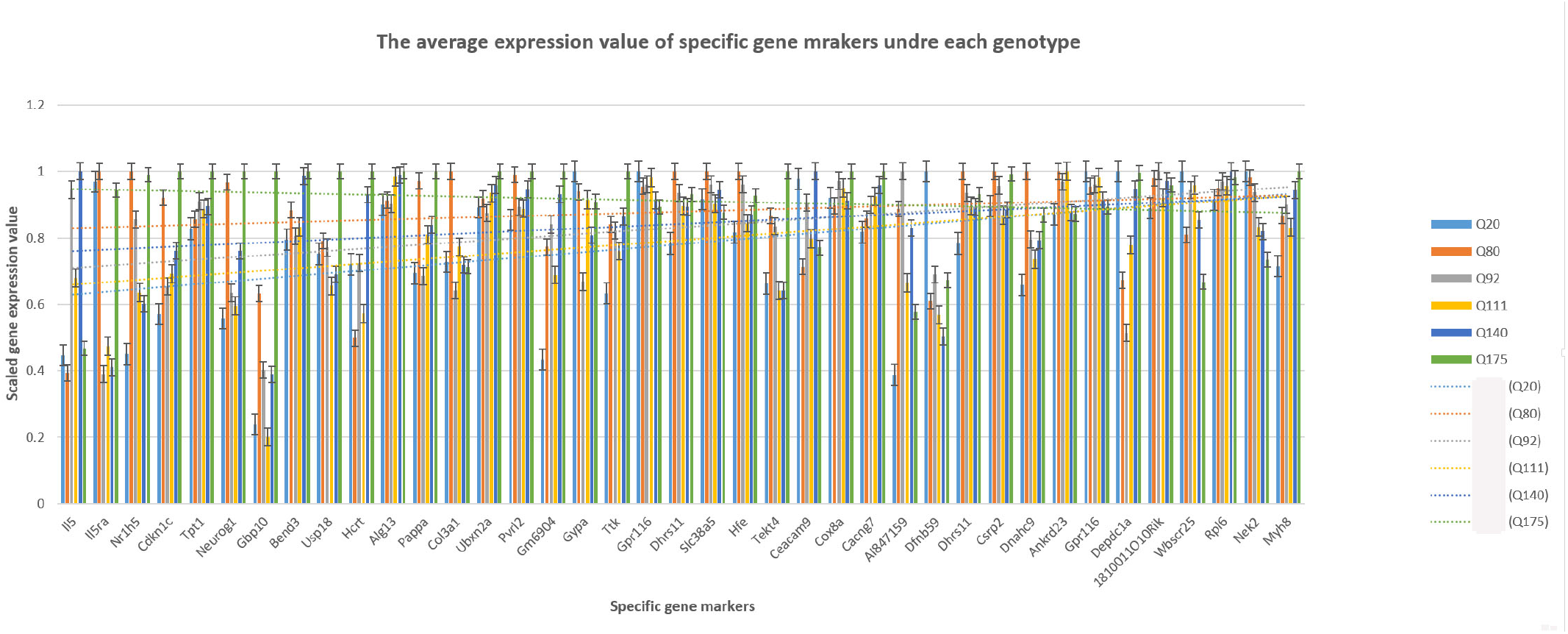
Scaled gene expression of the 39 marker genes with different genotypes.

## IV. Conclusion

Gene expression is characterized by spatiotemporal specificity, and genes often express in multi-tissues and different types of cells at different. It is hard to clearly describe the molecular process under complex clinical phenotypes. To further investigate molecular mechanisms that correlated to the repeat of CAG in Htt, we designed this research and k-means based unsupervised feature selection method to prioritize poly Q-related biomarkers. In our study, different biomarker genes were selected out for different phenotype. We further analysis the reason for the disease severity of different genotypes. Five human essential genes, Tpt1, Neurog1, Alg13, Col3a1 and Ttk are prioritized to be seriously affected by the mutation Htt gene. Besides, immune related pathways are related to the disease.

## ACKNOWLEDGMENT

This work was supported by grants from National Key R D Program of China (No. 2017YFC0909200), National Natural Science Foundation of China (No. 81671328 and No. 81971292), Program for Professor of Special Appointment (Eastern Scholar) at Shanghai Institutions of Higher Learning (No. 1610000043).

## Notes

This work was supported by grants from National Natural Science Foundation of China (No. 61381671328), Program for Professor of Special Appointment (Eastern Scholar) at Shanghai Institutions of Higher Learning (No. 1610000043), Innovation Research Plan supported by Shanghai Municipal Education Commission (ZXWF082101).

### Competing Interest Statement

The authors have declared no competing interest.

## References

[1] S. E. Browne, A. C. Bowling, U. Macgarvey, et al, “ Oxidative damage and metabolic dysfunction in Huntington’s disease: selective vulnerability of the basal ganglia,” Annals of Neurology, vol. 41, num. 5, pp. 646–653, 2010.

[2] T. Seredenina, R. Luthi-Carter, “What have we learned from gene expression profiles in Huntington’s disease?” Neurobiology of Disease, vol. 45, num. 1, 2012.

[3] N. F. Bence, R. M. Sampat, R. R. Kopito, “Impairment of the Ubiquitin-Proteasome System by Protein Aggregation,” Science, vol 292, pp. 1552–1555, 2001.

[4] N. S. Wexler, etal, “Venezuelan kindreds reveal that genetic and environmental factors modulate Huntington’s disease age of onset,” Proceedings of the National Academy of Sciences, USA, vol. 101, num. 10, pp. 3498–3503, 2004.

[5] A. Rosenblatt, R. R. Brinkman, K. Y. Liang, etal, “Familial influence on age of onset among siblings with Huntington disease,” American Journal of Medical Genetics, vol. 105, num. 5, pp. 399–403, 2001.

[6] M. Bjrkqvist, E. J. Wild, S. J. Tabrizi, “Harnessing immune alterations in neurodegenerative diseases,” Neuron, vol. 64, num. 1, pp. 21–24, 2009.

[7] E. Wild, M. Bjorkqvist, S. J. Tabrizi, “Immune markers for Huntington’s disease?” Expert Rev. Neurother., vol. 8, pp. 1779–1781, 2008.

[8] M. Bjorkqvist, E. J. Wild, J. Thiele, et al, “A novel pathogenic pathway of immune activation detectable before clinical onset in Huntingtons disease,” J.Exp. Med., vol. 205, pp. 1869–1877, 2008.

[9] M. Bjorkqvist, E. J. Wild, J. Thiele, et al, “A novel pathogenic pathway of immune activation detectable before clinical onset in Huntingtons disease,” J.Exp. Med., vol. 205, num. 205, pp. 1869–1877, 2008.

[10] Q. Fang, et al, “Brain-specific proteins decline inthe cerebrospinal fluid of humans with Huntington disease,” Mol. Cell. Proteomics, vol. 8, pp. 451–466, 2009.

[11] S. Hochreiter, J. Schmidhuber, “Long short-term memory,”Neural Computation, vol. 9, no. 8, pp: 1735–1780, 1997.

[12] Y. LeCun, “Generalization and network design strategies,” In: Connectionism in perspective. Citeseer, 1989.

[13] Y. LeCun, L. Bottou, Y. Bengio, P. Haffner, “Gradient-based learning applied to document recognition,” Proceedings of the IEEE, vol. 86, no. 11, pp:2278–2324, 1998.

[14] S. C. IoffeS, “Batch normalization: accelerating deep network training by reducing internal covariate shift,” In: International conference on machine learning,2015.

[15] Y. Bengio, P. Simard, P. Frasconi, “Learning long-term dependencies with gradient descentis difficult,” IEEE Trans NeuralNetw, vol. 5, no. 2, pp:157–166, 2002.

[16] R. Pascanu, T. Mikolov, Y. Bengio, “On the difficulty of training recurrent neural networks,” In:International conference on machine learning, PP:1310–1318, 2013.

[17] P. Shannon, A. Markiel, O. Ozier, N.S. Baliga, J.T. Wang, D. Ramage, et al., Cytoscape: a software environment for integrated models of biomolecular interaction networks., Genome research, 2003, vol. 13, no. 11, pp. 2498504.

[18] R. Oughtred, C. Stark, B.-J. Breitkreutz, J. Rust, L. Boucher, C. Chang, et al., The BioGRID interaction database: 2019 update., Nucleic Acids Research, 2019, vol. 47, no. D1, pp. D529D541

[19] F. Hong, R. Breitling, “A comparison of meta-analysis methods for detecting differentially expressed genes in microarray experiments,” Bioinformatics, vol. 24, no. 3, pp. 374–382, 2008.

[20] C. A. Ross, E. H. Aylward, E. J. Wild, et al., “Huntington disease: natural history, biomarkers and prospects for therapeutics,” Nature Reviews Neurology, vol. 10, no. 4, pp: 204–216, 2014.

[21] P. Langfelder, J. P. Cantle, D. Chatzopoulou, et al., “Integrated genomics and proteomics define huntingtin CAG length-dependent networks in mice,” Nature Neuroscience, vol. 19, no. 4, pp:623–633, 2016.

[22] K. Wang, M. Li, and H. Hakonarson, ANNOVAR: functional annotation of genetic variants from high-throughput sequencing data., Nucleic Acids Research, 2010, vol. 38, no. 16, pp. e164 e164.

